# Type VI secretion system killing by commensal *Neisseria* is influenced by the spatial dynamics of bacteria

**DOI:** 10.1101/2020.11.26.400259

**Authors:** Rafael Custodio, Rhian M. Ford, Cara J. Ellison, Guangyu Liu, Gerda Mickute, Christoph M. Tang, Rachel M. Exley

## Abstract

Type VI Secretion Systems (T6SS) are widespread in bacteria and can dictate the development and organisation of polymicrobial ecosystems by mediating contact dependent killing. In *Neisseria* species, including *Neisseria cinerea* a commensal of the human respiratory tract, interbacterial contacts are mediated by Type four pili (Tfp) which promote formation of aggregates and govern the spatial dynamics of growing *Neisseria* microcolonies. Here we show that *N. cinerea* expresses a plasmid-encoded T6SS that is active and can limit growth of related pathogens. We explored the impact of Tfp expression on *N. cinerea* T6SS-dependent killing and show that expression of Tfp by prey strains enhances their susceptibility to T6SS, by keeping them in close proximity of T6SS-wielding attacker strains. Our findings have important implications for understanding how spatial constraints during contact-dependent antagonism can shape the evolution of microbial communities.

## INTRODUCTION

The human microbiota is critical for the development of a healthy gastrointestinal immune system (Round and Mazmanian, 2009; Sommer and Bäckhed, 2013) and can also protect the host from invasion by pathogenic bacteria (Kamada et al., 2013). The microbes that carry out these important functions live as part of complex communities shaped by their fitness and ability to adapt to their environment, and which can be remodeled through mutualistic and antagonistic interactions (Garcia-Bayona and Comstock, 2018; Little et al., 2008; Nadell et al., 2016). Competition for niche and host-derived resources has therefore driven the evolution in bacteria of an array of mechanisms to suppress growth of, or kill neighbouring microbes. One mechanism, the Type VI Secretion System (T6SS), provides an effective strategy to eliminate competitors in a contact-dependent manner. The T6SS is a contractile, bacteriophage-like nanomachine that delivers toxins into the cytosol of target organisms (Cianfanelli et al., 2016; Ho et al., 2014). T6SS-associated effectors possess a broad range of activities, including nucleases (Koskiniemi et al., 2013; Ma et al., 2014; Pissaridou et al., 2018), phospholipases (Flaugnatti et al., 2016; Russell et al., 2013), peptidoglycan hydrolases (Whitney et al., 2013) and pore-forming proteins (Mariano et al., 2019); each effector is associated with a cognate immunity protein to prevent self-intoxication (Alcoforado Diniz et al., 2015; Unterweger et al., 2014). T6SSs have been best characterised in pathogenic bacteria, including *Pseudomonas*, *Vibrio*, *Salmonella* and *Shigella*, where its impact in pathogenesis and bacterial competition has been established *in vitro* and in some cases *in vivo* (Anderson et al., 2017; Sana et al., 2016). However, commensal bacteria also harbour T6SS, although how these systems combat pathogens has only been elucidated for *Bacteriodetes* in the intestinal tract (Russell et al., 2014); further studies are needed to gain a greater appreciation of how T6SS in commensals influence microbial communities and pathogens in other niches.

The human nasopharynx hosts a polymicrobial community (Kumpitsch et al., 2019; Marchesi et al., 2017; Ramos-Sevillano et al., 2019), which can include the obligate human pathobiont *Neisseria meningitidis*, as well as related but generally non-pathogenic, commensal *Neisseria* species (Diallo et al., 2016; Dorey et al., 2019; Gold et al., 1978; Knapp and Hook, 1988; Sheikhi et al., 2015). *In vivo* studies have demonstrated an inverse relationship between carriage of commensal *Neisseria lactamica* and *N. meningitidis* (Deasy et al., 2015), while *in vitro* studies have revealed that some commensal *Neisseria* demonstrate potentially antagonistic effects against their pathogenic relatives (Custodio et al., 2020; Kim et al., 2019). Commensal and pathogenic *Neisseria* species have also been shown to interact closely in mixed populations (Custodio et al., 2020; Higashi et al., 2011a). Social interactions among *Neisseria* are mediated by surface structures known as Type IV pili (Tfp). These filamentous organelles enable pathogenic *Neisseria* to adhere to host cells (Nassif et al., 1993; Virji et al., 1991), and are crucial for microbe-microbe interactions and the formation of bacterial aggregates and microcolonies (Helaine et al., 2007; Higashi et al., 2007). In addition, Tfp interactions can dictate bacterial positioning within a community; non-piliated strains have been shown to be excluded to the expanding edge of colonies growing on solid media (Oldewurtel et al., 2015; Zöllner et al., 2017) while heterogeneity in pili, for example through post translational modifications, can alter how cells integrate into micocolonies (Zöllner et al., 2017).

We recently demonstrated that the pathogen *N. meningitidis* closely interacts with commensal *Neisseria cinerea* on human epithelial cell surfaces in a Tfp-dependent manner (Custodio et al., 2020). Here, whole genome sequence analysis revealed that the *N. cinerea* isolate used in our studies encodes a T6SS. We provide the first description of a functional T6SS in *Neisseria* spp.. We show that the *N. cinerea* T6SS is encoded on a plasmid and antagonises pathogenic relatives, *N. meningitidis* and *Neisseria gonorrhoeae*. Moreover, we examined whether Tfp influence the competitiveness of microbes in response to T6SS-mediated antagonism and demonstrate that T6SS–mediated competition is facilitated by Tfp in bacterial communities.

## RESULTS

### N. cinerea *346T encodes a functional T6SS on a plasmid*

We identified a single locus in *N. cinerea* isolate CCUG346T (346T) (https://www.ccug.se/strain?id=346) that encodes homologues of all 13 components that are necessary for a functional T6SS (Cascales and Cambillau, 2012), including genes predicted to encode canonical T6SS components Hcp and VgrG (**Figure 1A** - **table supplement 1**). We used T6SS-effector prediction software tools (Li et al., 2015) to search for putative effectors. In total we identified six putative effector and immunity genes, termed *Nte* and *Nti* for <u>N</u>eisseria <u>T</u>6SS <u>e</u>ffector/<u>i</u>mmunity, respectively. All Ntes contain a conserved Rhs domain, frequently associated with polymorphic toxins (Busby et al., 2013), and a variable C-terminal region. Nte1 contains an N-terminal PAAR motif, which can associate with the VgrG tip of T6SS (Shneider et al., 2013) and C-terminal phospholipase domain (cd00618). Nte2 also contains an N-terminal PAAR domain and has a predicted RNase domain (pfam15606) in its C-terminal region, while Nte3 is a putative HNH endonuclease (pfam14411). Nte4 contains a GIY-YIG nuclease domain (cd00719) and Nte5 is predicted to be an HNH/endo VII nuclease (pfam14412), with Nte6 predicted to contain an HNHc endonuclease active site (cd00085).

**Figure 1.**
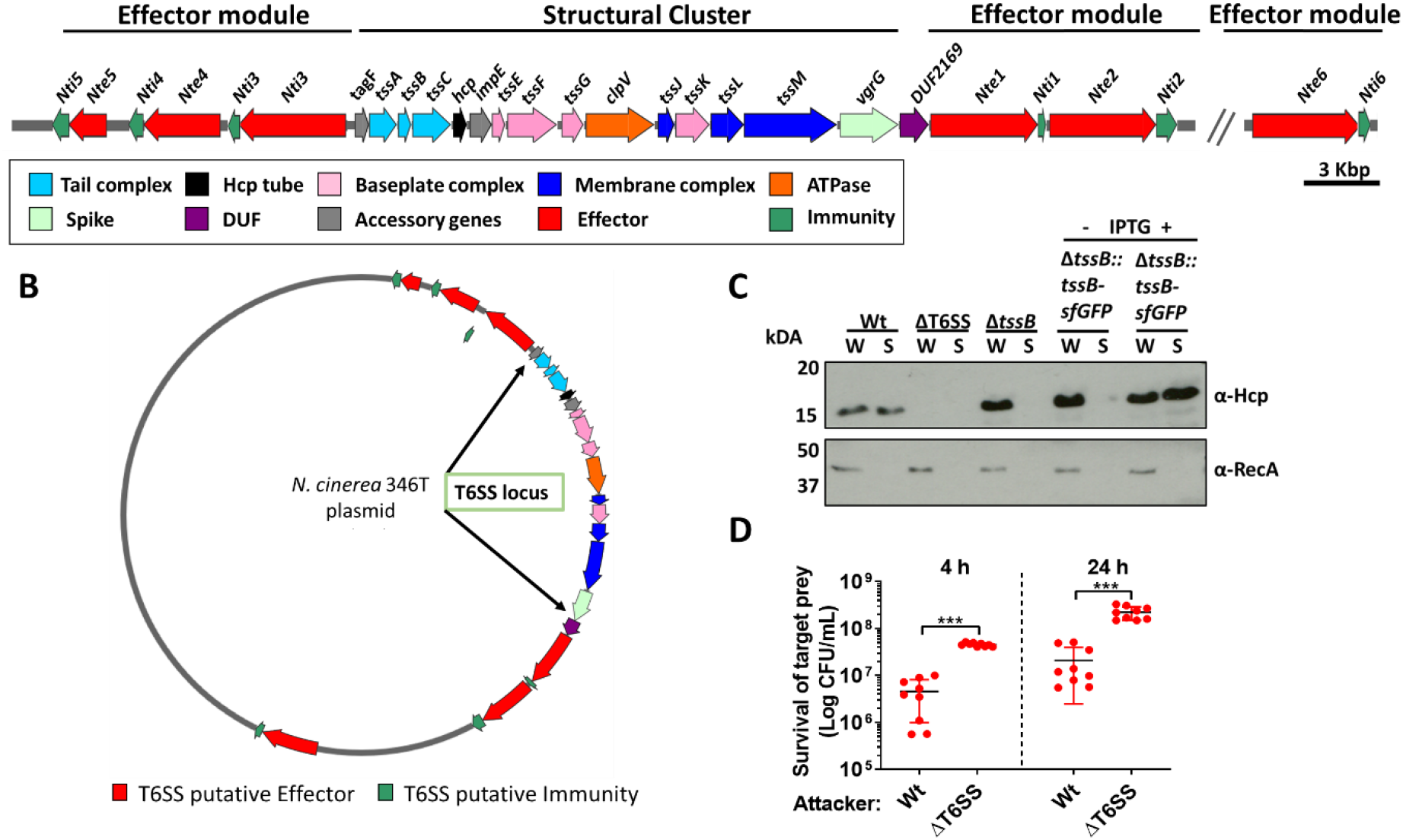
*N. cinerea* expresses a functional T6SS. **(A)**, Schematic representation of T6SS genes in *N. cinerea* 346T. Canonical *tss* nomenclature was used for genes in the T6SS cluster. **(B)** Map of the T6SS-associated genes encoded by the *N. cinerea* 346T plasmid. **(C)** Expression and secretion of Hcp by wild-type *N. cinerea* 346T (Wt) and the T6SS mutant (ΔT6SS). Hcp protein was detected in the whole cell lysates (W) and supernatants (S) by western blot analysis. For strain Δ*tssB*::*tssB*sfGFP, bacteria were grown in the presence (+) or absence (−) of 1 mM IPTG; molecular weight marker shown in kDa. RecA is only detected in whole cell lysates. **(D)** Survival of the prey, *N. cinerea* 27178A, after 4 and 24 co-incubation with wild-type *N. cinerea* 346T or the T6SS mutant (ΔT6SS) at 10:1 ratio, attacker:prey. The mean ± SD of three independent experiments are shown: ***p < 0.0001 using unpaired two-tailed Student’s t-test.

Of note, all predicted T6SS-related *orf*s and Nte/Ntis in *N. cinerea* 346T were found to be encoded on a 108,141 bp plasmid, revealed by PacBio sequencing, and confirmed by PCR and sequencing. Nte/Nti 1-5 are encoded adjacent to the structural genes cluster, with Nte6/Nti6 encoded elsewhere in the plasmid (**Figure 1B** - **figure supplement 1**). Thus, our analysis reveals that the human commensal *N. cinerea* harbours a plasmid-borne T6SS together with six putative effector-immunity pairs.

Contraction of T6SSs leads to Hcp secretion, a hallmark of a functional T6SS (Cascales and Cambillau, 2012). Therefore, to establish whether the *N. cinerea* T6SS is functional, we assessed Hcp levels in whole cell lysates and supernatants from wild-type *N. cinerea*, a ΔT6SS mutant lacking 10 core genes including *hcp,* and a Δ*tssB* mutant. As expected, Hcp was detected in both fractions from the wild-type strain but not in cell lysates or supernatants from the ΔT6SS mutant (**Figure 1C**). Importantly, Hcp was present in cell lysates from the Δ*tssB* mutant, but not detected in cell supernatants, consistent with TssB being a component of the T6SS-tail-sheath required for contraction (Brackmann et al., 2018). Hcp secretion was restored by complementation of the Δ*tssB* mutant by chromosomal expression of TssB with a C-terminal sfGFP fusion (Δ*tssB*::*tssB*-sfGFP) (**Figure 1C**).

Next, we performed competition assays between *N. cinerea* 346T or the ΔT6SS mutant against *N. cinerea* 27178A which lacks a T6SS and Nte/Nti pairs identified in *N. cinerea* 346T. The survival of *N. cinerea* 27278A was reduced by around an order of magnitude following incubation with *N. cinerea* 346T compared with the ΔT6SS mutant (**Figure 1D**, p < 0.0001 and p < 0.0001, respectively), confirming that the *N. cinerea* 346T T6SS is active during inter-bacterial competition.

### *Dynamic behaviour of the* Neisseria *T6SS in the presence of prey cells*

We further analysed the activity of the T6SS by visualising assembly and contraction in *N. cinerea* 346TΔ*tssB*::*tssB*-sfGFP; this strain exhibits comparable T6SS killing as wild-type *N. cinerea* 346T (**figure supplement 2**). Time-lapse microscopy revealed dynamic T6SS foci inside bacteria, with structures extending/contracting over seconds (**Figure 2A** - **movie supplement 1**) consistent with T6SS activity(Gerc et al., 2015; Ringel et al., 2017). To further confirm T6SS activity, we deleted the gene encoding the TssM homologue in strain 346TΔ*tssB*::*tssB*-sfGFP, abolishing T6SS activity (**Figure 2B**) and confirmed that in the absence of TssM, fluorescent structures were rarely seen (< 5% of cells in the Δ*tssM* background, compared with > 60% in the strain expressing TssM; **Figure 2C** - **movie supplement 2**).

**Figure 2.**
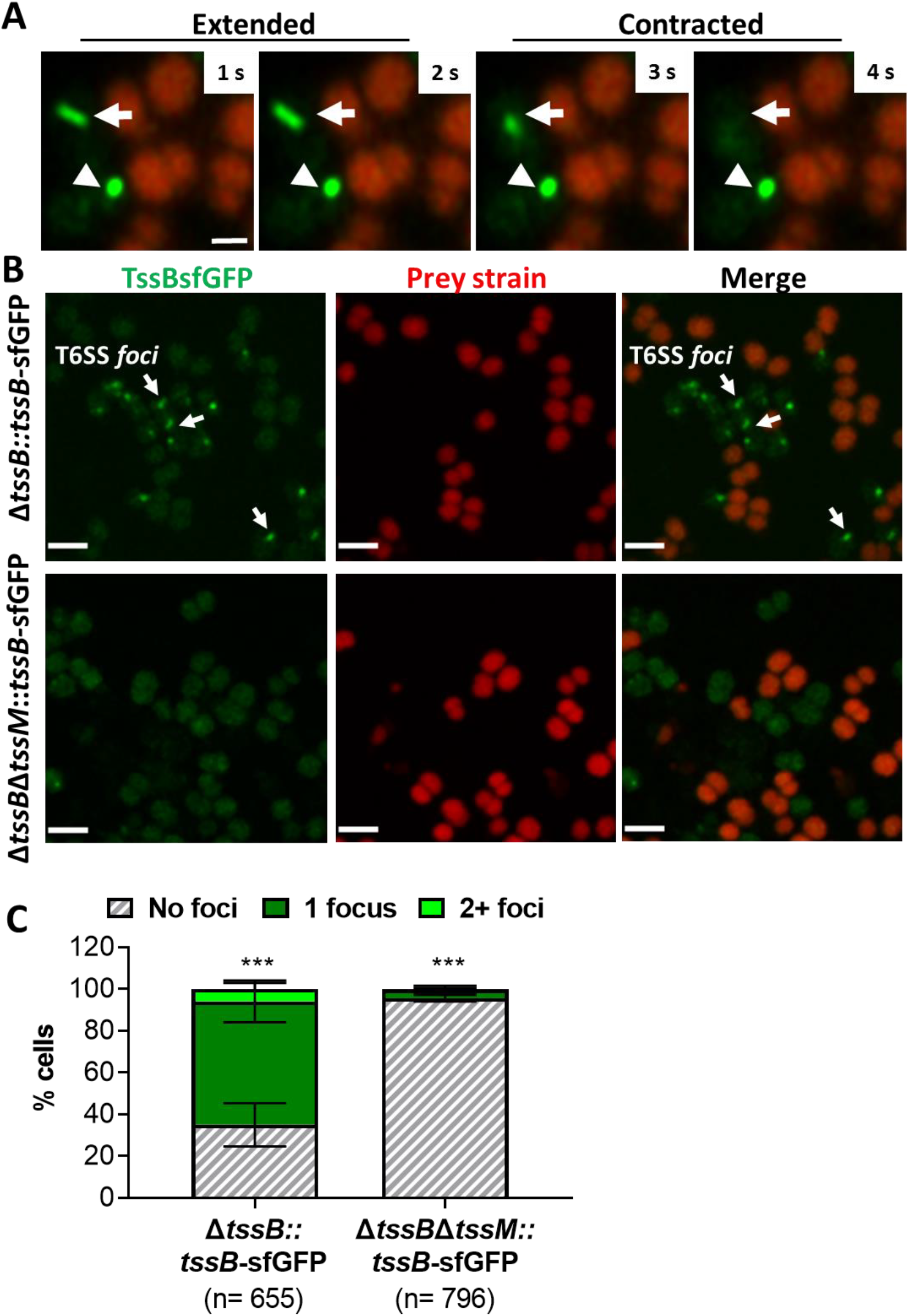
Visualisation of T6SS activity in *N. cinerea.* **(A)** Assembly and contraction of the T6SS in *N. cinerea*; white arrows indicate contracting T6SSs. Time-lapse images of *N. cinerea* 346TΔ*tssB*::*tssB*sfGFP (green) and prey *N. cinerea* 27178A_ *sfCherry* (red); the arrowhead shows a non-dynamic focus, scale bar, 1 μm. See also Movie supplement 1. **(B)**Representative images of *N. cinerea* strains with the TssB::sfGFP fusion with (upper panels) or without (lower panels) TssM. Loss of fluorescent foci upon deletion of *tssM* indicates that foci correspond to active T6SS. The scale bar represents 2 μm. **(C)**Quantification of TssB-sfGFP foci in different strains. T6SS foci were quantified using ‘analyse particle’ (Fiji) followed by manual inspection. For each strain, at least two images from gel pads were obtained on two independent occasions. Percentage of cells with 0, 1, or 2+ foci are shown and n = number of cells analysed. Data shown are mean ± SD of two independent experiments: ***p<0.0001 using two-way ANOVA test for multiple comparison. See also Movie supplement 2.

Finally, we examined whether T6SS assembly induces lysis of prey cells. We imaged *N. cinerea* 346TΔ*tssB*::*tssB*-sfGFP with *N. cinerea* 27178 expressing sfCherry on gel pads with SYTOX Blue as an indicator of target cell permeability (Ringel et al., 2017). Interestingly, we detected increased SYTOX staining of prey cells immediately adjacent to predator bacteria displaying T6SS activity (**Figure 3** - **movie supplement 3**), indicating that *N. cinerea* T6SS induces cell damage and lysis of its prey.

**Figure 3.**
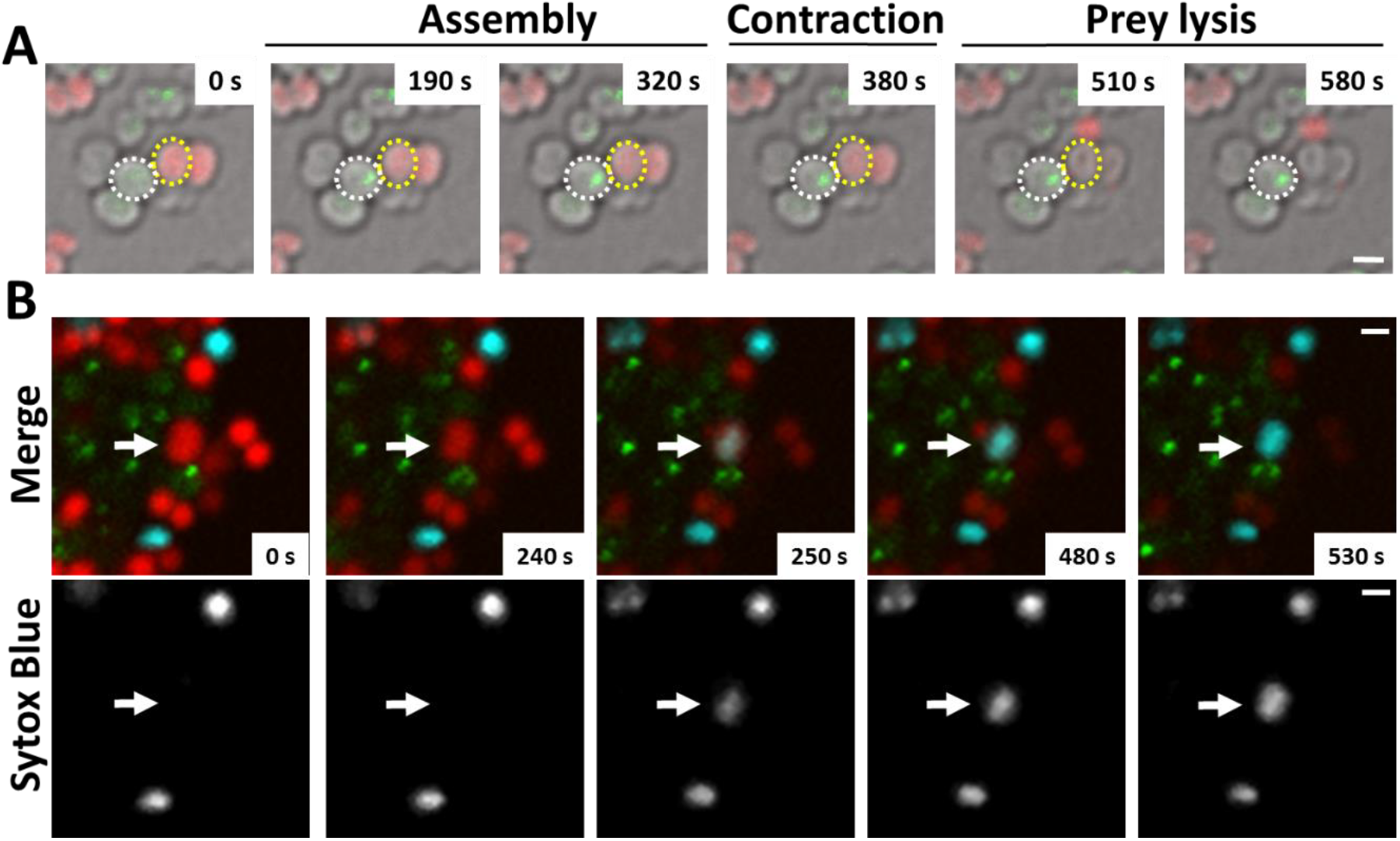
*N. cinerea* T6SS induces lysis in prey bacteria. **(A)** Assembly of T6SS and prey lysis. Time-lapse series of merged images with phase contrast, *N. cinerea* 346T Δ*TssB*+*TssB*sfGFP (green), and *N. cinerea* 27178A sfCherry (red); scale bar, 1 μm. **(B)** Top row shows merged images of GFP (green, indicating T6SS assembly/contraction), mCherry (red, prey strain), and SYTOX Blue (cyan, showing membrane permeabilisation) channels. The bottom row arrows highlight a prey cell losing membrane integrity (increase in SYTOX Blue staining inside cells) arrows. Representative image from two biological repeats. Scale bars represent 1 μm. See also Movie supplement 3.

### N. cinerea *T6SS effectors are functional toxin/immunity pairs*

To characterise the T6SS effector:immunity pairs, we expressed each Nte alone or with its corresponding Nti using an inducible expression plasmid in *E. coli*. We were only able to clone wild-type Nte6 in presence of its immunity protein, so Nte6^R1300S^ was used to analyse toxicity of this effector. In addition, as Nte1 encodes a predicted phospholipase that should be active against cell membranes (Flaugnatti et al., 2016), we targeted the putative phospholipase domain of Nte1 to the periplasm by fusing it to the PelB signal sequence (Singh et al., 2013); cytoplasmic expression of the Nte1 phospholipase domain does not inhibit bacterial growth (**figure supplement 3**). All Ntes are toxic, with their expression leading to decreased viability and reduced optical density (OD) of *E. coli* cultures compared to empty vector controls; toxicity was counteracted by co-expression of the corresponding Nti (**Figure 4A-F**). Overall, these findings provide evidence that all six Nte/Ntis are effector-immunity proteins.

**Figure 4.**
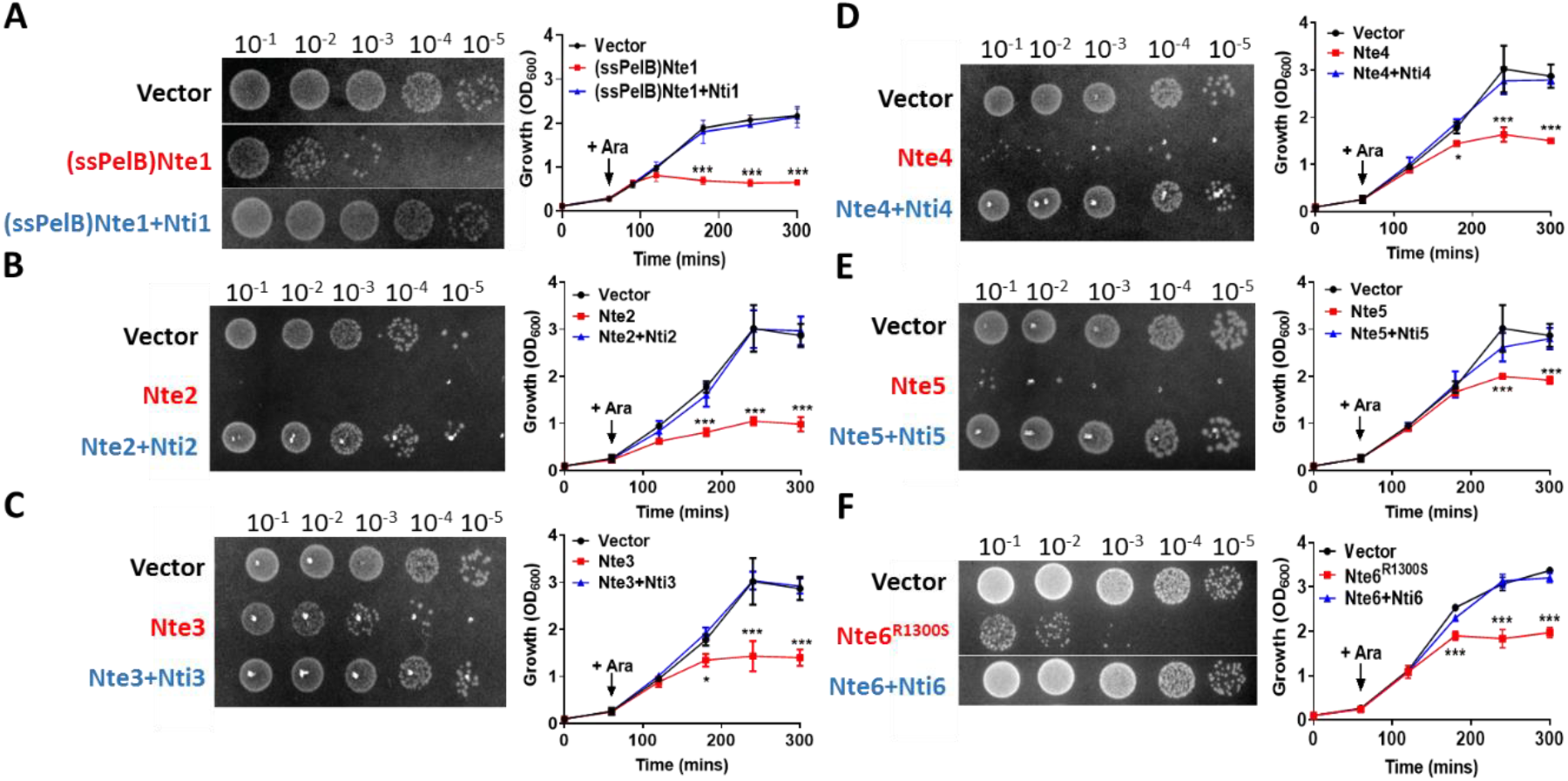
Putative *N. cinerea* T6SS effectors are toxic in *E. coli.* **(A)** Arabinose (Ara) induced expression of T6SS effector Nte1 in periplasm of *E. coli* leads to reduction in CFU and OD at 600 nm (OD_600_). Co-expression of putative immunity Nti1 restores growth to levels of strain with empty vector (pBAD33). **(B)-(E)** Cytoplasmic expression of putative effectors Nte2-5 without cognate immunity reduces growth and survival of *E. coli.* **(F)** Expression of Nte6^R1300S^ reduces viability and growth when expressed in *E. coli*. Expression of Nti6 with Nte6 does not impact growth. In (A)-(F) number of CFU at 120 mins post induction are shown. Data shown are the mean ± SD of three independent experiments: NS, not significant, ***p < 0.0001, *p < 0.05 using two-way ANOVA test for multiple comparison. Images of colonies for Nte1 and Nte6 are composite as strains were spotted to different areas of the same plates.

### *Commensal* Neisseria *T6SS kills human pathogens*

We next investigated whether commensal *N. cinerea* can deploy T6SS to antagonise the related pathogenic species, *N. meningitidis* and *N. gonorrhoeae*. We performed competition assays with three *N. meningitidis* strains (belonging to different lineages and expressing different polysaccharide capsules *i.e.* serogroup B or C), and a strain of *N. gonorrhoeae*. *N. cinerea* 346T caused between a 50 to 100-fold decrease in survival of the meningococcus compared with the ΔT6SS strain, irrespective of lineage or serogroup (**Figure 5A**) and an approximately 5-fold reduction is survival of the gonococcus (**Figure 5B**). We also investigated whether the meningococcal capsule protects against T6SS assault. Using a capsule-null strain (Δ*siaD*) in competition assays with wild-type *N. cinerea* 346T or the T6SS mutant, we found there was an approximately 5-fold reduction in survival of the Δ*siaD* mutant compared to the isogenic wild-type strain (**Figure 5C**). Therefore, the meningococcal capsule protects bacteria against T6SS attack.

**Figure 5:**
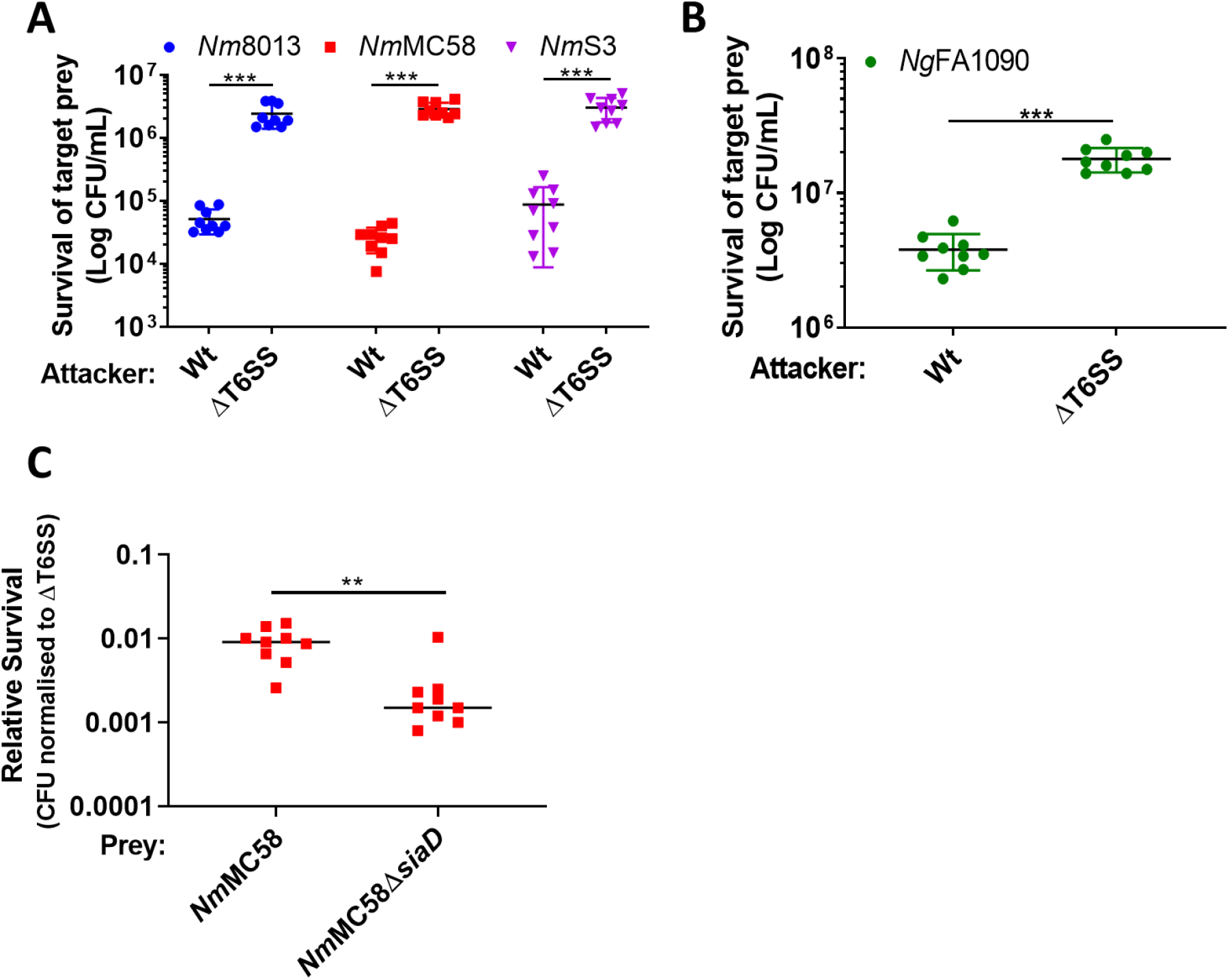
*N. cinerea* T6SS is active against pathogenic *N. meningitidis* and *N. gonorrhoeae*. **(A)** Recovery of wild-type *N. meningitidis* (*Nm*8013, *Nm*MC58, *Nm*S3) after 4 h co-incubation with *N. cinerea* 346T wild-type (Wt) or the T6SS mutant (ΔT6SS) at a 100:1 attacker:prey ratio. **(B)** Recovery of wild-type *N. gonorrhoeae* (FA1090) after 4 h co-incubation with *N. cinerea* 346T wild-type (Wt) or the T6SS mutant (ΔT6SS) at a 10:1 attacker:prey ratio, attacker:prey. **(C)** Unencapsulated *N. meningitidis* (*Nm*MC58Δ*siaD*) is more susceptible to T6SS-mediated killing than wild-type *N. meningitidis*. Recovery of *Nm*MC58 or the capsule-null mutant (*Nm*MC58Δ*siaD*) after 4 h co-culture with *N. cinerea* 346T (Wt) or a T6SS-deficient mutant (ΔT6SS) at ratio of 100:1, attacker:prey. Relative survival is defined as the fold change in recovery of *N. meningitidis* following incubation with wild-type *N. cinerea* compared to *N. cinerea* ΔT6SS. Data shown are the mean ± SD of three independent experiments: NS, not significant, ***p < 0.0001, **p < 0.001 using unpaired two-tailed Student’s t-test for pairwise comparison (B and C) or one-way ANOVA test for multiple comparison (A).

### Spatial segregation driven by Type IV pili dictates prey survival against T6SS assault

Despite the potency of T6SS in *Neisseria* warfare, this nanomachine operates when bacteria are in close proximity, so we hypothesised that Tfp, which are critical for the formation of *Neisseria* microcolonies and organisation of bacterial communities (Higashi et al., 2007; Mairey et al., 2006; Oldewurtel et al., 2015; Zöllner et al., 2017), could influence T6SS-mediated antagonism. To test this we constructed fluorophore expressing, piliated and non-piliated ‘prey’ strains (*i.e*. sfCherry-expressing 346TΔ*nte*/*i3-5* which is sensitive to T6SS-mediated attack (**figure supplement 4**). Prey strains were mixed with piliated attacker strain *N. cinerea* 346T expressing sfGFP at a 1:1 ratio on solid media, and the spatiotemporal dynamics of bacterial growth examined by time-lapse stereo microscopy over 24 h, while the relative proportion of each strain was analysed by flow cytometry at 24 h (**figure supplement 5**). As expected based on previous observations of Tfp-mediated cell sorting in *Neisseria* (Oldewurtel et al., 2015; Zöllner et al., 2017), the non-piliated prey strain (346TΔ*nte*/*i3-5*Δ*pilE1*/*2*_*sfCherry*; red) segregates to the periphery of the colony, in this location the prey strain escapes T6SS-mediated assault and dominates the expanding colony (**Figure 6A** - **movie supplement 4 and figure supplement 5**). In contrast, when the prey is piliated, pilus-mediated cell interactions prevent displacement of cells to the expanding front (Oldewurtel et al., 2015; Ponisch et al., 2018; Zöllner et al., 2017), so the susceptible strain (Tfp-expressing 346TΔ*nte*/*i3-5_sfCherry* Tfp+, red) is outcompeted by the T6SS+ strain (Tfp-expressing 346T_*sfGfp* Tfp+, green) (**Figure 6B** - **movie supplement 5 and figure supplement 5**). When both strains are piliated and immune to T6SS attack, there is no dominance of either strain (**Figure 6C** - **movie supplement 6 and figure supplement 5**). Assessment of the relative recovery of piliated and non-piliated prey in competition assays also supported the observation that the piliation status of the prey impacts survival against T6SS (**Figure 6D)**. These results highlight that, Tfp influence the outcome of T6SS-mediated antagonism through structuring and partitioning bacteria in mixed microcolonies.

**Figure 6:**
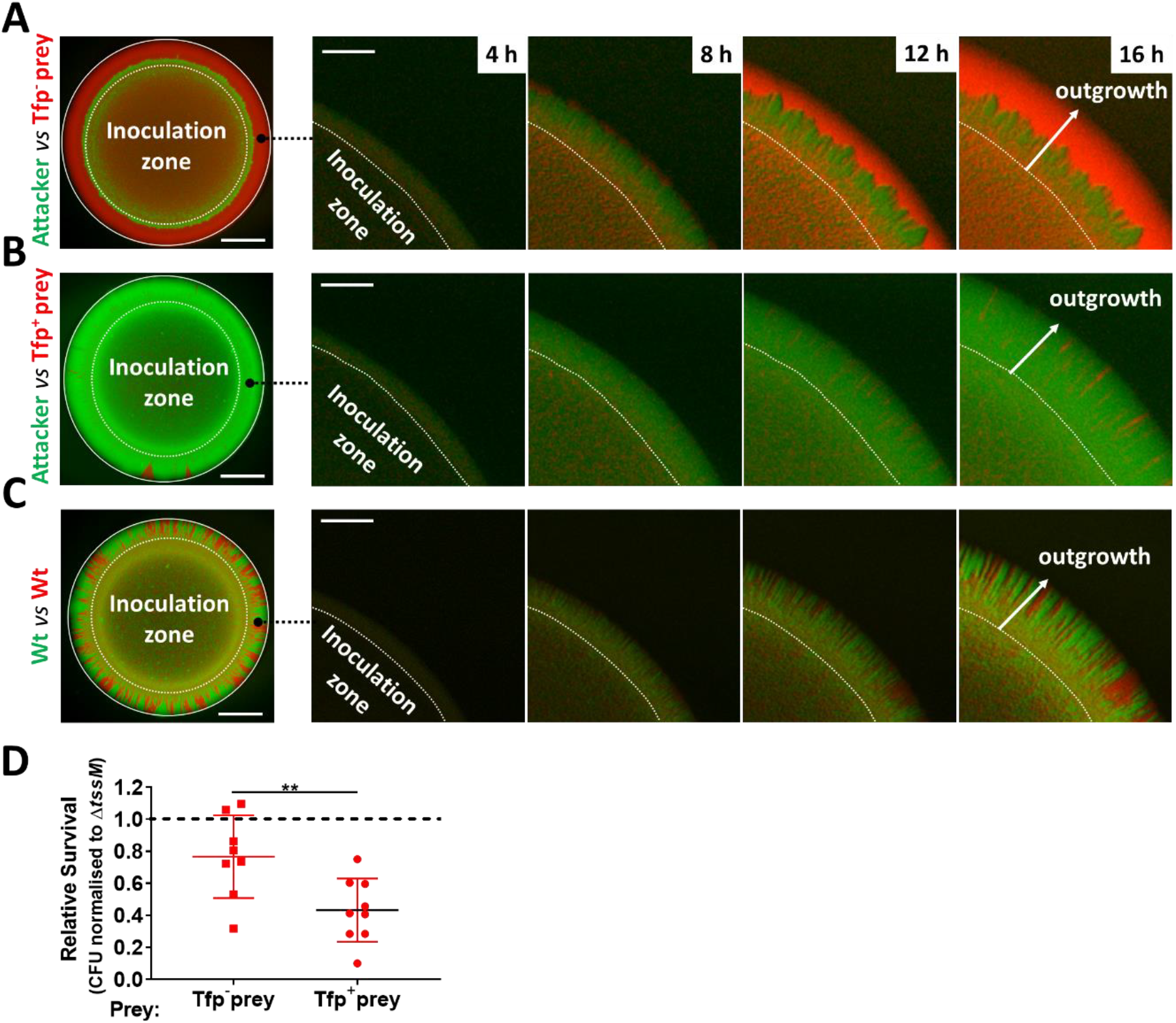
Predator and prey piliation promotes T6SS killing. **(A)** Fluorescence microscopy images taken at specific times after inoculation of mixed (1:1 ratio) bacterial colonies. A T6SS-susceptible, non-piliated prey strain (346TΔ*nte*/*i3*-*5*Δ*pilE1*/*2_sfCherry*, red) migrates to the expanding edge of the colony over time, segregating from the T6SS+ attacker strain (*N. cinerea* 346T*_gfp*, green) and dominating the expanding population. **(B)** The same susceptible prey strain but expressing pili does not segregate, and after 24h is outcompeted by the piliated T6SS+ attacker. **(C)** The non-T6SS-susceptible piliated prey strain (346T*_sfCherry*, red) and piliated attacker strain (346T_sf*Gfp*, green) do not segregate, but due to immunity against T6SS attack, no dominance is observed. Images of colonies are representative of three independent experiments. Scale bar, 500 μm. Expanding colony edge images are stills at indicated times from time-lapse imaging performed on one occasion. Scale bar 100 μm. Movies supplement 4-6 show expanding colonies in A-C, respectively. **(D)** The influence of piliation on T6SS killing. Recovery of non-piliated and piliated prey strains after 24 h co-culture with *N. cinerea* 346T (Wt) and a *tssM*-deficient mutant (Δ*tssM*) at ratio of 10:1, attacker:prey. Relative survival is defined as the fold change in recovery of prey following incubation with wild-type attacker *N. cinerea* compared to *N. cinerea* Δ*tssM*. Data shown are the mean ± SD of three independent experiments: **p < 0.01 using unpaired two-tailed Student’s t-test for pairwise comparison.

## DISCUSSION

Here we identified a T6SS in a commensal *Neisseria* spp. which can kill T6SS-deficient *N. cinerea* isolates and the related pathogens, *N. meningitidis*, with which it shares an ecological niche (Knapp and Hook, 1988), and *N. gonorrhoeae*. Of note, the *N. cinerea* T6SS is encoded on a large plasmid, with structural genes for the single T6SS apparatus clustered in one locus, similar to other T6SS (Anderson et al., 2017; Liaw et al., 2019; Sana et al., 2016). Effectors Nte1 to 5 are also encoded in the same locus, but with Nte6 encoded elsewhere on the plasmid. To date, plasmid encoded T6SS have only been described in *Campylobacter* species (Marasini and Fakhr, 2016), with this plasmid T6SS mobilised *via* conjugation (Marasini et al., 2020). Although other small plasmids have been reported in *N. cinerea* (Knapp et al., 1984; Roberts, 1989) and *N. cinerea* can be a recipient of *N. gonorrhoeae* plasmids (Genco et al., 1984), it is not yet known whether T6SS plasmids are widespread among *Neisseria*, or whether the plasmid can be mobilised by conjugation or transformation. Interestingly, in *Acinetobacter baylyi*, T6SS induced prey cell lysis contributes to acquisition of plasmids from target cells (Ringel et al., 2017). Therefore, it will be interesting to see whether other *Neisseria* species with T6SS genes (Marri et al., 2010) harbour T6SS-expressing plasmids.

Examination of *N. cinerea* T6SS activity revealed several interesting features. Microscopy demonstrated that T6SS attack (tit-for-tat) is not required to provoke firing of the system. Instead, the T6SS appears to be constitutively active in *N. cinerea* (**Figure 2**). Furthermore, the system is capable of inducing lysis of prey bacteria (**Figure 3**). The consequences of T6SS attack are determined by the repertoire and activities of effectors, and their site of delivery. Many different effector activities have been proposed including lipases, peptidoglycan hydrolases, metalloproteases and nucleases (Lewis et al., 2019). Effector activities can result in target cell lysis to varying degrees (Ringel et al., 2017; Smith et al., 2020). Of the six Ntes we identified, lysis could be mediated by Nte1 which harbours a putative phospholipase domain in the C-terminus. Alternatively, a combination of effectors might be needed to elicit prey lysis.

Polysaccharide capsules are largely thought to provide bacteria with a strategy for evading host immune killing (Lewis and Ram, 2014). Here, we found that the meningococcal capsule has an alternative role in defence against other bacteria. Meningococcal strains lacking a capsule were at a significant disadvantage in the face of a T6SS-expressing competitor implicating this surface polysaccharide in protection against T6SS assault. Similar findings have been reported for other bacteria; for example the extracellular polysaccharide of *V. cholerae* and the colonic acid capsule of *E. coli* confer defence against T6SS attack (Hersch et al., 2020; Toska et al., 2018). One potential mechanism is that the capsule sterically impairs the ability of the T6SS to penetrate the target cell membrane, and/or inhibits access of T6SS effectors to their cellular targets. Interestingly, recent genetic evidence indicates that some commensal *Neisseria* species also have capacity to produce polysaccharide capsules (Clemence et al., 2018), which might also confer a survival advantage in mixed populations that include strains expressing T6SS.

Most bacteria exist within complex polymicrobial communities in which the spatial and temporal dynamics of proliferation and death has a major effect on their fitness and survival (Nadell et al., 2016). While structured complex microbial societies can benefit all their members (Gabrilska and Rumbaugh, 2015; Wolcott et al., 2013), antagonistic neighbours, especially those deploying contact-dependent killing mechanisms, can disrupt communities. Although T6SS-mediated killing can be advantageous to a producing strain during bacterial competition, this requires intimate association with its prey (MacIntyre et al., 2010; Russell et al., 2014). Thus, one way for susceptible bacteria to evade T6SS killing is to avoid direct contact with attacking cells (Borenstein et al., 2015; Smith et al., 2020). In *Neisseria*, the Tfp is a key mediator of interbacterial and interspecies interactions (Custodio et al., 2020; Higashi et al., 2011b) and pilus-mediated interactions influence the spatial structure of a growing community (Oldewurtel et al., 2015; Zöllner et al., 2017). In *N. gonorrhoeae,* non-piliated bacteria segregate to the expanding front of the colony and Tfp-mediated spatial reorganisation can allow bacteria to avoid external stresses or strains competing for resources (Oldewurtel et al., 2015; Zöllner et al., 2017). We predicted that this would be especially relevant in the context of T6SS mediated antagonism, where physical exclusion driven by Tfp-loss or modification, which may occur naturally in a polymicrobial environment and is an established phenomenon in pathogenic *Neisseria* (Hagblom et al., 1985; Helm and Seifert, 2010), could be an effective strategy to evade and survive an antagonistic interaction, while pilus-mediated interactions might be less favourable for a susceptible prey. Our results demonstrate that piliation of the predator and prey strains of *N. cinerea* led to dominance of the T6SS-expressing strain, indicating that *N. cinerea* Tfp amplify T6SS-mediated competition. It is noteworthy that many bacteria (*e.g. Pseudomonas aeruginosa*, *Vibrio cholerae*, *Acinetobacter baumannii*, enteropathogenic *E. coli*) that employ T6SS for inter-bacterial competition also express Tfp. Therefore, our findings are of broad relevance for the impact of contact dependent killing, and further emphasise how precise spatial relationships can have profound effects on how antagonistic and mutualistic factors combine to influence the development of microbial communities.

## MATERIALS AND METHODS

### Bacterial strains and growth

Bacterial strains used in this study are shown in **Table Supplement 2**. *Neisseria* spp. were grown on Brain Heart Infusion (BHI, Oxoid) agar with 5% defibrinated horse blood or in BHI broth at 37°C with 5% CO_2_ or GC-medium supplemented with 1.5% base agar (w/v) and 1% Vitox (v/v; Oxoid). GW-medium (Wade and Graver, 2007) was used for *N. cinerea* microscopy experiments. *E. coli* was grown on LB (Lennox Broth base, Invitrogen) agar or in liquid LB at 37°C with shaking. Antibiotics were added at the following concentrations: for *E. coli*, carbenicillin (carb) 100 μg/ml, kanamycin (kan) 50 μg/ml, and chloramphenicol (cm) 20 μg/ml; for *Neisseria* spp. kan 75 μg/ml, spectinomycin (spec) 65 μg/ml, erythromycin (ery) 15 μg/ml, and polymyxin B (pmB) 10 μg/ml.

### DNA Isolation and whole-genome sequencing (WGS)

Genomic DNA was extracted using the Wizard Genomic Kit (Promega), and sequenced by PacBio (Earlham Institute, Norwich) using single-molecule real-time (SMRT) technology; reads were assembled *de novo* with HGAP3(Chin et al., 2013).

### *Construction of* N. cinerea *mutants*

Primers used in this study are listed in **Table supplement 3**. Target genes were replaced with antibiotic cassettes as previously (Wörmann et al., 2016). Constructs were assembled into pUC19 by Gibson Assembly (New England Biolabs), and hosted in *Escherichia coli* DH5α. Plasmids were linearised with *Sca*I, and gel extracted relevant linearised fragments used to transform *N. cinerea*; transformants were checked by PCR and sequencing. Complementation or chromosomal insertion of genes encoding fluorophores was achieved using pNCC1-Spec, a spectinomycin-resistant derivative of pNCC1(Wörmann et al., 2016). For visualisation of T6SS-sheaths, *sfgfp* was cloned in-frame with *tssB* and a short linker (encoding 3×Ala 3×Gly) by Gibson Assembly (New England Biolabs) into pNCC1-Spec to allow IPTG-inducible expression of TssB-sfGFP. PCR was performed using Herculase II (Agilent) or Q5 High-fidelity DNA Polymerase (New England Biolabs).

### Analysis of effector/immunity activity in E. coli

Putative effector coding sequences with or without cognate immunity gene were amplified by PCR from *N. cinerea* 346T gDNA and either assembled by Gibson Assembly (NEB) into pBAD33 or, for Nte1 with or without addition of the PelB signal sequence, cloned in to pBAD33 using XbaI / SphI restriction enzyme sites. Plasmids were transformed into *E. coli* DH5α and verified by sequencing (Source Bioscience). For assessment of toxicity, strains with recombinant or empty pBAD33 plasmids were grown overnight in LB supplemented with 0.8% glucose (w/v), then diluted to an OD_600_ of 0.1 and incubated for 1 hour at 180 rpm and 37°C; bacteria were pelleted and resuspended in LB with arabinose (0.8% w/v) to induce expression and incubated at 37°C, 180 rpm for a further 4 h. The OD_600_ and CFU/ml of cultures were determined; aliquots were diluted and plated to media containing 0.8% glucose at relevant time points up to 5 h.

### Hcp protein expression, purification and antibody generation

Codon optimised *hcp* was synthesised with a sequence encoding an N-terminal 6x His Tag and a 3C protease cleavage site, and flanked by *Nco*I and *Xho*I restriction sites (ThermoFisher). The fragment was ligated into *Nco*I and *Xho*I sites in pET28a (Novagen) using QuickStick T4 DNA Ligase (Bioline) and transformed into *E. coli* B834. Bacteria were grown at 37°C, 150 rpm to an OD_600_ of 1.0, and expression of 6xHis-3C-Hcp was induced with 1 mM IPTG for 24 h at 16°C. Cells were resuspended in Buffer A (50 mM Tris-HCl buffer pH 7.5, 10 mM Imidazole, 500 mM NaCl, 1 mM DTT) containing protease inhibitors, 1 mg/mL lysozyme and 100 μg/mL DNase then subsequently homogenised with an EmulsiFlex-C5 (Avestin). Lysed cells were ultracentrifuged and the cleared supernatant loaded onto a Ni Sepharose 6 Fast Flow His Trap column (GE Healthcare) equilibrated with Buffer A. The column was washed with Buffer A, then Buffer B (50 mM Tris-HCl buffer pH 7.5, 35 mM Imidazole, 500 mM NaCl, 1 mM DTT) before elution with 10 mL of Buffer C (50 mM Tris-HCl buffer pH 7.5, 300 mM Imidazole, 150 mM NaCl, 1 mM DTT). The eluate was incubated with the HRV-3C protease (Sigma) then applied to a Ni Sepharose column. The eluate containing protease and cleaved protein was concentrated using Amicon Ultra 10,000 MWCO (Millipore), then passed through a Superdex-200 column (GE Healthcare, Buckinghamshire, UK). Fractions were analysed by SDS-PAGE and Coomassie blue staining, and those with Hcp pooled and used to generate polyclonal antibodies (EuroGentec).

### Hcp secretion assay

Bacteria were grown in BHI broth for 4-5 h then harvested and lysed in an equal volume of SDS-PAGE lysis buffer (500 mM Tris-HCl [pH 6.8], 5% SDS, 15% glycerol, 0.5% bromophenol blue containing 100 mM β-mercaptoethanol); supernatants were filtered (0.22 μm pore, Millipore) and proteins precipitated with 20% (v/v) trichloroacetic acid. Hcp was detected by Western blot with anti-Hcp (1:10,000 dilution) and goat anti-rabbit IgG–HRP (1:5000, sc-2004; Santa Cruz). Anti-RecA (1:5000 dilution, ab63797; Abcam) followed by goat anti-rabbit IgG–HRP and detection with ECL detection Reagent (GE Healthcare) or Coomassie blue staining were used as loading controls.

### Live cell imaging of T6SS activity

Bacteria were grown overnight on BHI agar, resuspended in PBS and 20 μl spotted onto fresh BHI agar containing 1 mM IPTG and incubated for 4 h at 37°C. After incubation, 500 μl of 10^9^ cfu/mL bacterial suspension of attacker was mixed with the prey strain at a 1:1 ratio. Cells were harvested by centrifugation for 3 min at 6000 rpm, resuspended in 100 μL of PBS or GW media and 2 μl spotted on 1% agarose pads (for T6SS dynamics) or GW media with 0.1 mM IPTG and 0.5 μM SYTOX™Blue (Thermo Fisher Scientific) for assessment of prey permeability. Fluorescence microscopy image sequences were acquired within 20-30 minutes of sample preparation with an inverted Zeiss 880 Airyscan microscope equipped with Plan-Apochromat 63×/1.4-NA oil lens and fitted with a climate chamber mounted around the objective to perform the imaging at 37°C with 5% CO_2_. Automated images were collected at 1 sec, 10 sec or 1 min intervals and processed with Fiji (Schindelin et al., 2012). Background noise was reduced using the “Despeckle” filter. The XY drift was corrected using StackReg with “Rigid Body” transformation (Thévenaz et al., 1998). Experiments and imaging were performed on at least two independent occasions.

### Quantitative competition assays

Strains grown overnight on BHI agar were resuspended in PBS and diluted to 10^9^ CFU/mL, mixed at the indicated ratio, then 20 μl spotted onto BHI agar in triplicate and incubated at 37°C with 5% CO_2_. At specific time-points, entire spots were harvested and resuspended in 1 mL of PBS. The cellular suspension was then serially diluted in PBS and aliquots spotted onto selective media. Colonies were counted after ~16 h incubation at 37 °C with 5% CO_2_. Experiments were performed on at least three independent occasions. For different prey analysis relative survival was defined as the fold change in recovery of prey following incubation with wild-type attacker *N. cinerea* compared to a T6SS-deficient *N. cinerea*.

### Competition assays assessed by Fluorescence microscopy and flow cytometry

Bacteria were prepared and grown as described for quantitative competition assays except using 1 μl spot volume and spotted onto GC-medium supplemented with 0.5% base agar (w/v) and 1% Vitox (v/v; Oxoid). At various time points expanding colonies were imaged using a M125C stereo microscope equipped with a DFC310FX digital camera (Leica Microsystems) and images processed with Fiji. Images were imported using “Image Sequence” and corrected with StackReg as described. For flow cytometry, colonies were harvested, fixed with 4% PFA for 20 min then washed with PBS. Samples were analysed using a Cytoflex LX (Beckman Coulter), and at least 10^4^ events recorded. Fluorescence, forward and side scatter data were collected to distinguish between debris and bacteria. Results were analysed by calculating the number of events positive for either GFP or Cherry signal in FlowJo v10 software (Becton Dickinson Company). Events that were negative for fluorescence or positive for both markers were also plotted. Flow cytometry analysis was performed on two independent occasions. Stereo microscopy analysis was performed on three independent occasions with technical duplicates each time.

### Statistical analyses

Graphpad Prism7 software (San Diego, CA) was used for statistical analysis. We used One-way/two-way ANOVA with Tukey post hoc testing for multiple comparisons and unpaired two-tailed Student’s t-test for pairwise comparisons. In all cases, *p* < 0.05 was considered statistically significant.

## ACKNOWLEDGEMENTS

We thank members of the Foster group (Oxford) especially Daniel Unterweger (now at the University of Kiel) for advice and assistance with microscopy as well as Alan Wainman of the SWDSP Bioimaging facility. We are grateful to M. Basler (Basel) for valuable advice, and to Meningitis Now for funding. Work in CMT’s lab is supported by a Wellcome Trust Investigator award (102908/Z/13/Z).

## SUPPLEMENTARY FIGURES

**Figure supplement 1.**
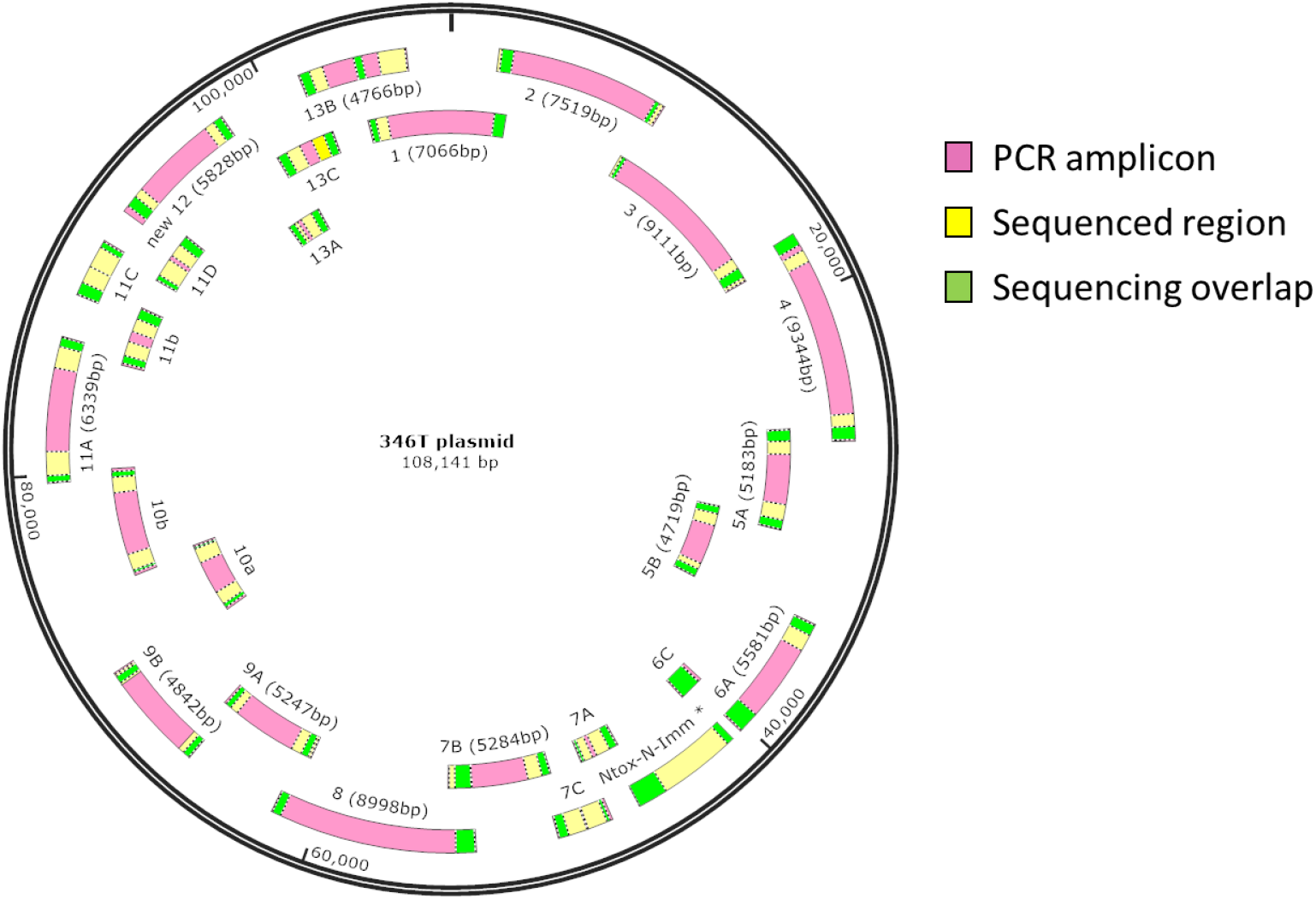
The *N. cinerea* 346T T6SS is encoded on a plasmid. Overlapping PCR and sequencing confirms extra-chromosomally closed circular DNA fragment. A total of 25 PCR fragments (pink bars) were amplified from *N. cinerea* 346T gDNA to confirm the plasmid predicted by PacBio whole-genome sequencing. Yellow shows regions which were sequenced, and green indicates the overlapping amplified regions.

**Figure supplement 2.**
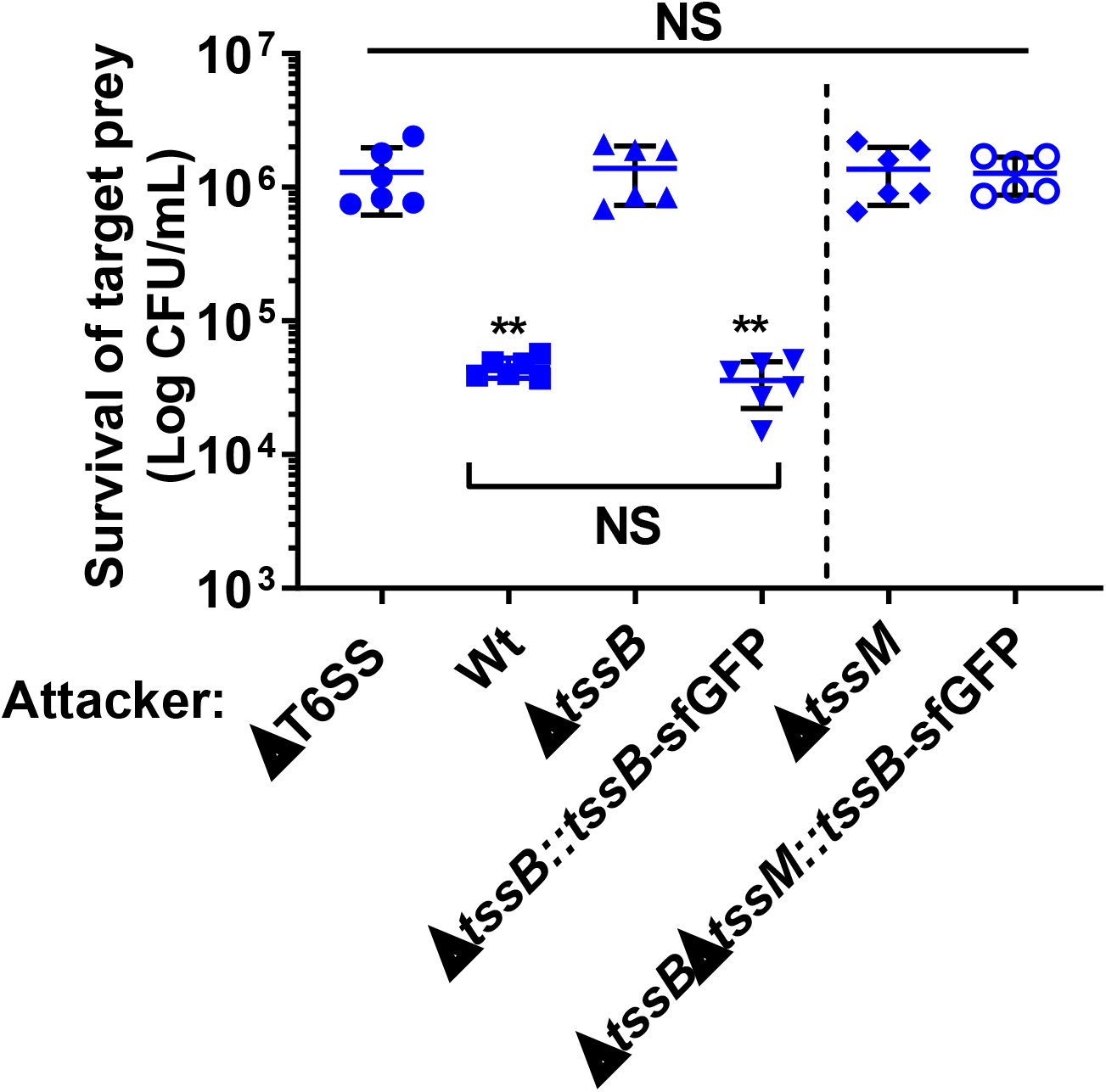
*N. cinerea* T6SS with a TssB C-terminal sfGFP fusion is functional and activity is lost upon deletion of *tssM.* Competition assay measuring the recovery of prey (*N. meningitidis* 8013) after 4 h co-incubation with wild-type *N. cinerea* 346T (Wt) and specified mutants at ratio of 100:1 (attacker:prey). Data shown are the mean ± SD of two independent experiments: NS, not significant, **p < 0.005 using one-way ANOVA test for multiple comparisons.

**Figure supplement 3.**
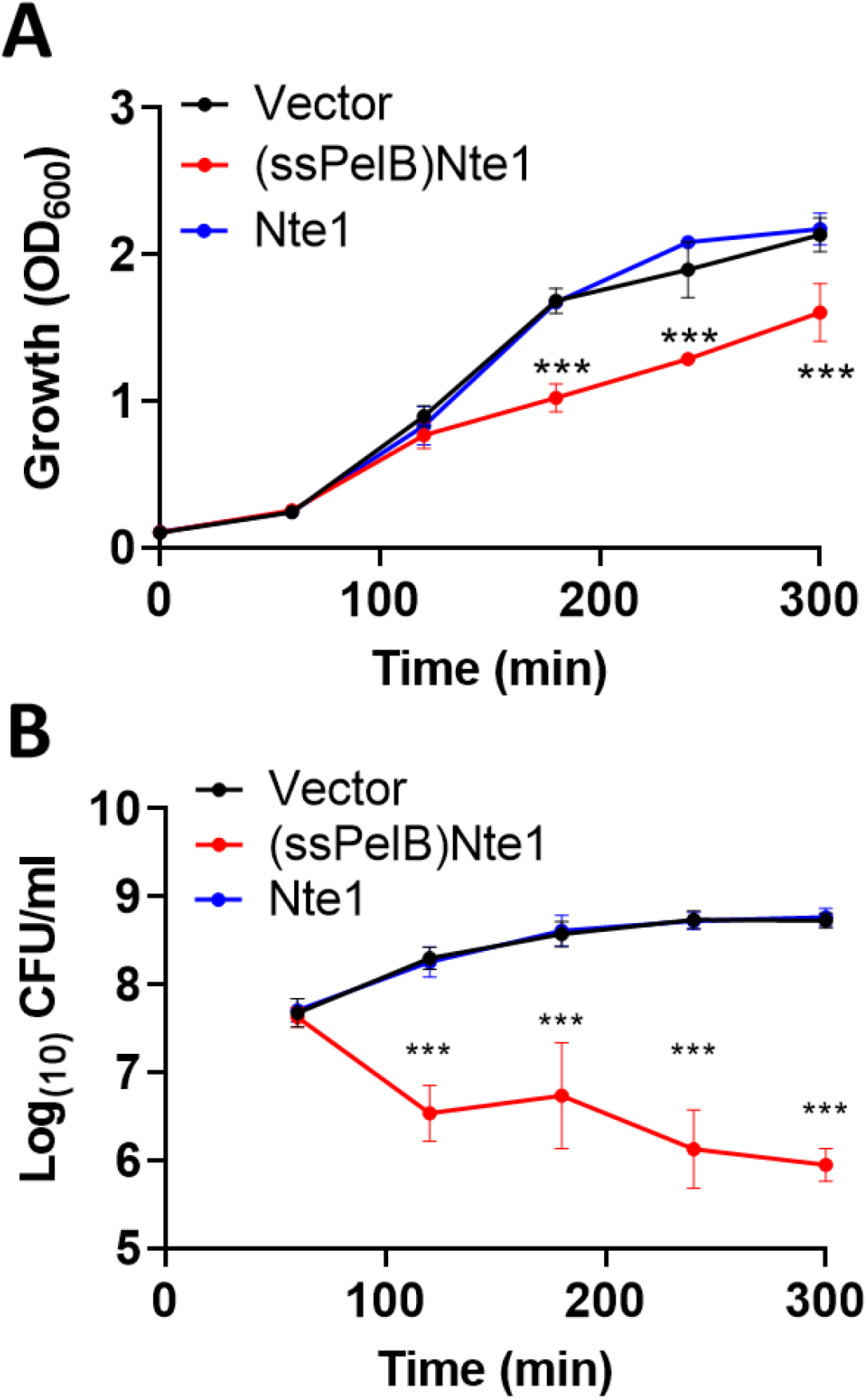
*N cinerea* putative T6SS effector Nte1 requires a PelB signal sequence for toxicity in *E. coli.* (**A**) Toxicity of Nte1 with or without the PelB signal sequence (ssPelB) following expression in *E. coli*. Expression was induced with L-arabinose at 60 mins and bacterial growth was monitored by measuring the OD_600_ of cultures. A reduction on OD600 was only observed when Nte1 was expressed with ssPelB. (**B**) Samples collected after addition of arabinose were plated to media containing glucose to repress toxin expression and enumerated. Data, shown are mean ± SD of three independent experiments: NS, not significant, ***p < 0.0001, *p < 0.05 using two-way ANOVA test for multiple comparison.

**Figure supplement 4.**
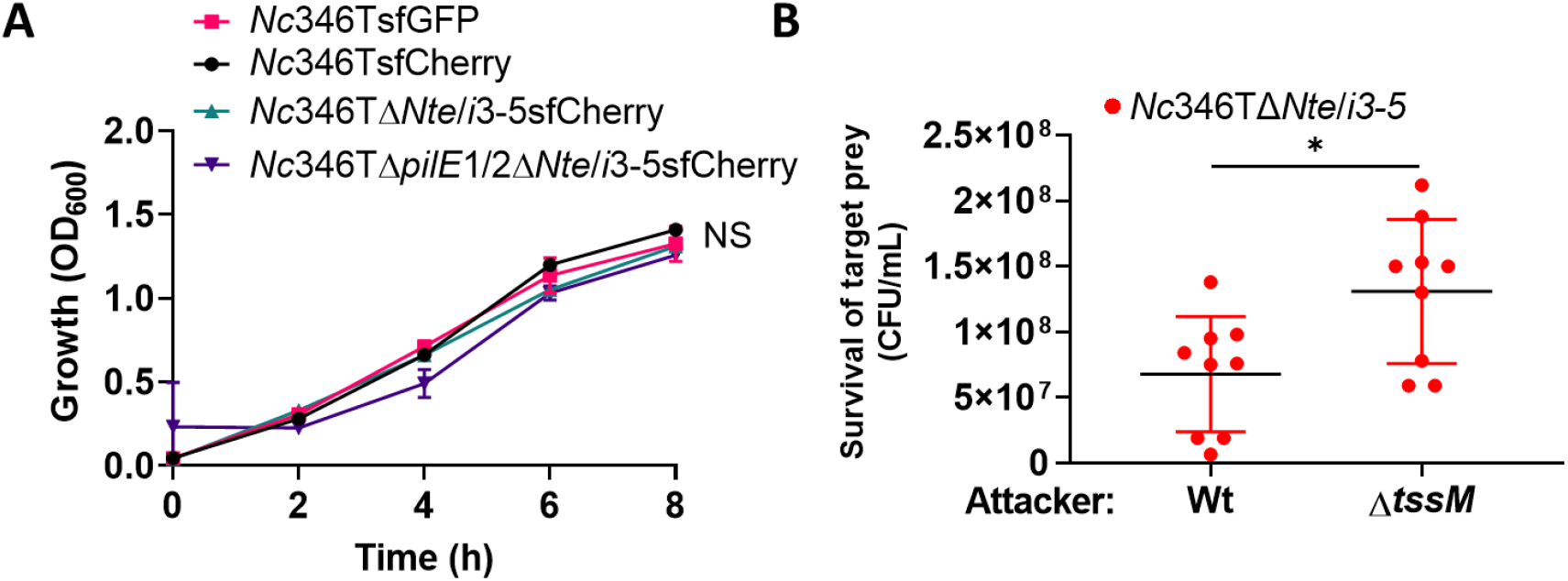
*N. cinerea* 346TΔ*nte*/*i3*-*5* has comparable growth to parent strain and is susceptible to T6SS-killing by wild-type *N. cinerea* 346T. (**A**) *N. cinerea* 346T strains were grown in liquid BHI media for 8 h at 37 ̊C with 5% CO_2_. Growth was monitored by measuring the OD_600_ of cultures. Data are representative of two independent experiments: NS, not significant, using two-way ANOVA test for multiple comparison. (**B**) Competition assay measuring the recovery of the indicated *N. cinerea* 346TΔ*Nte*/*i3*-*5* mutant strain after 4 h of co-incubation wild-type *N. cinerea* 346T (Wt) or a *tssM*-deficient mutant (Δ*tssM*) at ratio of 10:1 (attacker: prey). Data shown are the mean ± SD of three independent experiments performed in triplicate: *p < 0.05 using unpaired two-tailed Student’s t-test.

**Figure supplement 5.**
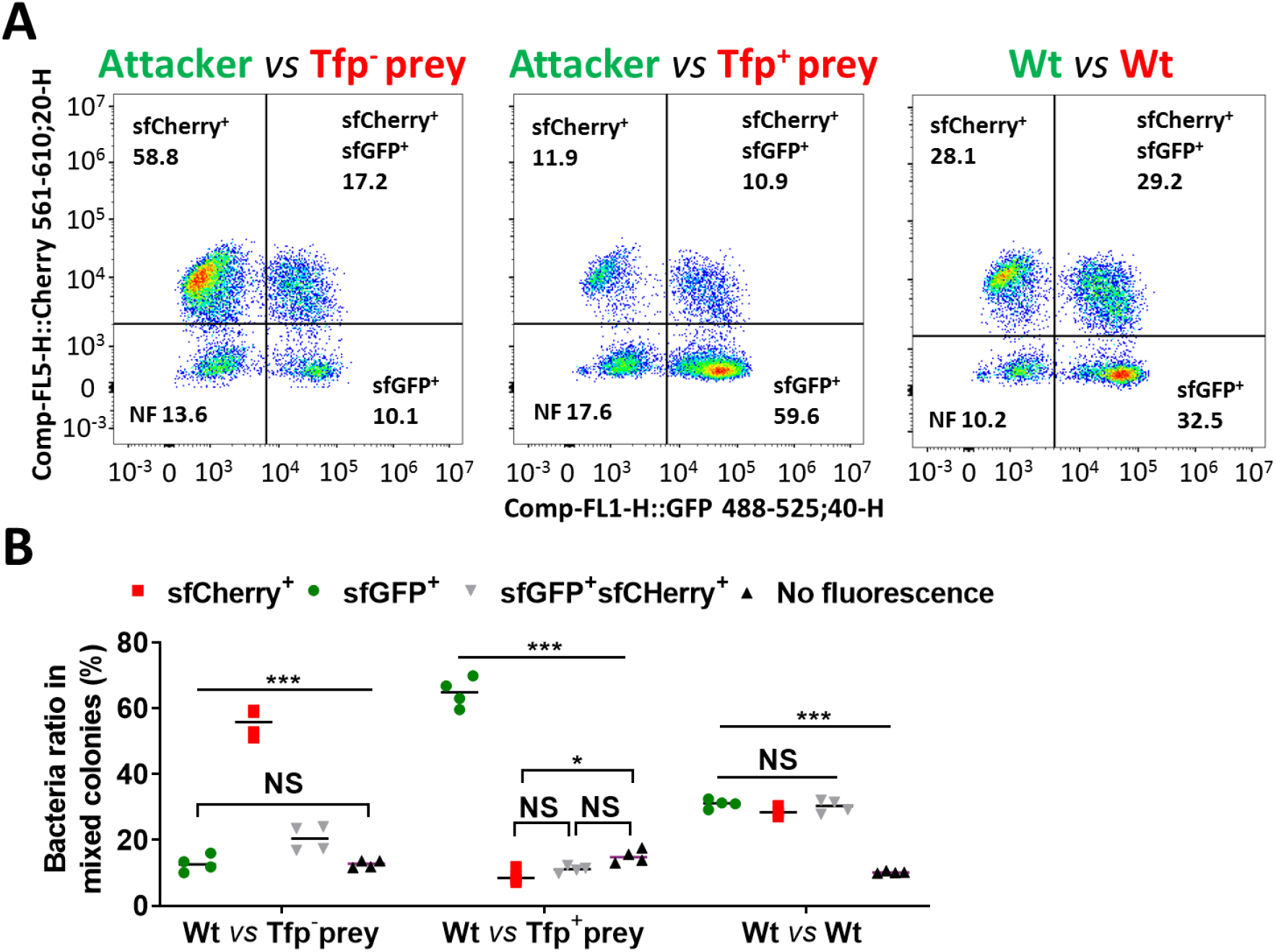
Flow cytometry analysis of relative proportion of piliated and non-piliated prey in mixed colonies with T6SS-expressing piliated attacker strain. **(A)** Total bacteria in mixed colonies grown on agar plates were collected, fixed and analysed by Flow cytometry. Plots reveal shift in the dominant population after 24h growth. Plots are representative of two independent experiments with two technical repeats. **(B)** Histograms of data shown in panel a. When non-piliated, after 24 h the prey strain (red dots) dominates the population, despite being susceptible to T6SS-mediated attack (red prey 346TΔ*Nte*/*i3-5*Δ*pilE1*/*2*_*sfCherry* 56±4%, green attacker 346T_*gfp* 13±3%; *p* < 0.0001). Piliation of the prey reduces it to less than 10% of the population, and the attacker cells constitute the majority of the population (red prey 346TΔ*Nte*/*i3-5* 9±2%, green attacker 346T_*gfp* 65±5%, *p* < 0.0001). When the immunity of the prey is not compromised, piliated attacker and prey are detected in approximately equal proportions (red 346T_sfCherry 29±1%, green 346T_*gfp* 31±1%, *p* = 0.58. Data, shown are mean ± SD of two independent experiments: NS, not significant, * p < 0.05, ***p < 0.0001 using two-way ANOVA test for multiple comparison.

## SUPPLEMENTARY TABLES

**Table supplement 1.** Putative T6SS core components in *N. cinerea* 346T.

| Name | Protein Size (aa) | % aa (coverage) identity with *P. aeruginosa* PAO1 | COG | Putative Localisation | Predicted Function |
| --- | --- | --- | --- | --- | --- |
| **TssJ** | 215 | 35 (58) | COG3521 | Outer membrane | Membrane complex |
| **TssL** | 421 | 36 (91) | COG3455 | Inner membrane | Membrane complex |
| **TssM** | 1185 | 27 (93) | COG3523 | Inner membrane | Membrane complex |
| **TssK** | 447 | 37 (99) | COG3522 | Inner membrane | Baseplate complex |
| **TssF** | 640 | 33 (99) | COG3515 | Inner membrane | Baseplate complex |
| **TssG** | 346 | 37 (92) | COG3520 | Inner membrane | Baseplate complex |
| **TssE** | 170 | 37 (97) | COG3518 | Cytoplasmic | Baseplate complex |
| **TssA** | 355 | 28 (98) | COG3515 | Cytoplasmic | Tail complex |
| **TssB** | 172 | 70 (98) | COG3157 | Cytoplasmic | Tail complex (sheath) |
| **TssC** | 499 | 73 (97) | COG3517 | Cytoplasmic | Tail complex (sheath) |
| **ClpV** | 883 | 59 (98) | COG0542 | Cytoplasmic | ATPase |
| **Hcp** | 160 | 41 (100) | COG3157 | Cytoplasmic/Inner membrane | Tail complex (Hcp tube) |
| **VgrG** | 757 | 35 (99) | COG3501 | Inner membrane | Spike |

**Table supplement 2.** Bacterial strains used in this study.

| Strains or plasmids | Description | Ref/<br>Source |
| --- | --- | --- |
| ***Neisseria cinerea*** |  |  |
| CCUG346T (346T) | wild-type *N. cinerea* | Bennett et al., 2012 |
| CCUG27178A (27178A) | wild-type *N. cinerea* | Bennett et al., 2012 |
| 346T_sfGFP | 346T with chromosomally integrated *sfGfp*; Ery<sup>R</sup> | Wörmann et al., 2016 |
| 346T_sfCherry | 346T with chromosomally integrated *sfCherry*; Ery<sup>R</sup> | This study |
| 27178A_sfCherry | 27178 with chromosomally integrated *sfCherry*; Spec<sup>R</sup> | This study |
| 346TΔT6SS | 346T with insertion-deletion of *tssC – vgrG* region; Ery<sup>R</sup> | This study |
| 346TΔtssB | 346T with insertion-deletion of *tssB*; Ery<sup>R</sup> | This study |
| 346TΔtssB::tssBsfGFP | 346T with insertion-deletion of native *tssB* and ectopic chromosomal insertion of *tssB-sfGFP* fusion; Spec<sup>R</sup> Ery<sup>R</sup> | This study |
| 346TΔtssM | 346T with insertion-deletion of *tssM*; Tet<sup>R</sup> | This study |
| 346TΔtssBΔtssM::tssB-sfGFP | 346T with insertion-deletion of native *tssB* and *tssM* and ectopic chromosomal insertion of *tssB-sfGFP* fusion; Spec<sup>R</sup> Ery<sup>R</sup> Tet<sup>R</sup> | This study |
| 346TΔnte/i3-5_sfCherry | 346T with insertion-deletion of *nte/i3-5* region and ectopic chromosomal insertion of *sfCherry*; Spec<sup>R</sup> Ery<sup>R</sup> | This study |
| 346TΔnte/i3-5Δpile1/2_sfCherry | 346T with insertion-deletion of *nte/i3-5* region; ectopic chromosomal insertion of *sfCherry*; insertion-deletion of *pile1* and *pile2*; kan<sup>R</sup>, Spec<sup>R</sup> Ery<sup>R</sup> | This study |
| ***Neisseria meningitidis*** |  |  |
| 8013 | *N. meningitidis* wild-type | Rusniok et al., 2009 |
| MC58 | *N. meningitidis* wild-type | Tettelin et al., 2000 |
| S3 | *N. meningitidis* wild-type | Uria et al., 2008 |
| MC58ΔsiaD | deletion mutagenesis, *NEIS0051*; Kan<sup>R</sup> | Virji et al., 1995 |
| ***Neisseria gonorrhoeae*** |  |  |
| FA1090 pGCC4 | FA1090 with chromosomally integrated plasmid pGCC4; Ery<sup>R</sup> | Mehr and Seifert, 1997 |
| ***Escherichia coli*** |  |  |
| Dh5α | Cloning strain | Lab collection |
| Dh5α pNCC1-Spec | Dh5α with pNCC1<sup>2</sup> Spec<sup>R</sup> plasmid | This study |
| Dh5α pNCC1-Spec-sfGFP | Dh5α with pNCC1-Spec with sfGFP insert; | This study |
| Dh5α pNCC101-Spec-sfCherry | DH5α with plasmid pNCC101+sfCherry insert. Spec<sup>R</sup> | Lab collection |
| Dh5α pUC19::ΔtssB | DH5α with pUC19::ΔtssB deletion construct; Carb<sup>R</sup> Ery<sup>R</sup> | This study |
| Dh5α pUC19::ΔtssM | DH5α with pUC19::ΔtssM deletion construct; Carb<sup>R</sup> Tet<sup>R</sup> | This study |
| Dh5α pUC19::ΔT6SS | DH5α with pUC19::ΔtssC-vgrG locus deletion construct; Carb<sup>R</sup> Ery<sup>R</sup> | This study |
| Dh5α pUC19:: Δnte3Δnte4nNte5 | DH5α with pUC19::Δnte3Δnte4nNte5 region including respective immunity genes deletion construct; Carb<sup>R</sup> Ery<sup>R</sup> | This study |
| B834 pET28a | B834 with pET28a IPTG-inducible expression vector, Amp<sup>R</sup> | Lab collection |
| Dh5α pET28a-His-3C-Hcp | Dh5α with pET28a vector for IPTG inducible expression of Nc 346T Hcp with N-terminal cleavable HIS tag. Amp<sup>R</sup> | This study |
| B834 pET28a-His-3C-Hcp | B834 expression strain, with pET28a vector for IPTG inducible expression of Nc 346T Hcp with N-terminal cleavable HIS tag. Amp<sup>R</sup> | This study |
| Dh5α pBAD33 | Dh5α with pBAD33 vector for *Arabinose-inducible expression*, Cm<sup>R</sup> | Lab collection |
| Dh5α pBAD33::(ssPelB)Nte1-His | Dh5α with pBAD33 encoding *Nte1 with N-terminal PelB leader peptide and C-terminal his-tag under arabinose-inducible promoter control*; Cm<sup>R</sup> | This study |
| Dh5α pBAD33:: (ssPelB)Nte1+Nti1 | Dh5α with pBAD33 encoding *Nte1 with N-terminal PelB leader peptide and C-terminal his-tag plus Nti, under arabinose-inducible promoter control*; Cm<sup>R</sup> | This study |
| Dh5α pBAD33::Nte1-His | Dh5α with pBAD33 encoding *Nte1 with N-terminal his-tag under arabinose-inducible promoter control*; Cm<sup>R</sup> | This study |
| Dh5α pBAD33::Nte2 | Dh5α with pBAD33 encoding *Nte2 under arabinose-inducible promoter control*; Cm<sup>R</sup> | This study |
| Dh5α pBAD33::Nte2+Nti2 | Dh5α with pBAD33 encoding *Nte2+Nti2 under arabinose-inducible promoter control*; Cm<sup>R</sup> | This study |
| Dh5α pBAD33::Nte3 | Dh5α with pBAD33 encoding *Nte3 under arabinose-inducible promoter control*; Cm<sup>R</sup> | This study |
| Dh5α pBAD33::Nte3+Nti3 | Dh5α with pBAD33 encoding *Nte3+Nti3 under arabinose-inducible promoter control*; Cm<sup>R</sup> | This study |
| Dh5α pBAD33::Nte4 | Dh5α with pBAD33 encoding *Nte4 under arabinose-inducible promoter control*; Cm<sup>R</sup> | This study |
| Dh5α pBAD33::Nte4+Nti4 | Dh5α with pBAD33 encoding *Nte4+Nti4 under arabinose-inducible promoter control*; Cm<sup>R</sup> | This study |
| Dh5α pBAD33::Nte5 | Dh5α with pBAD33 encoding *Nte5 under arabinose-inducible promoter control*; Cm<sup>R</sup> | This study |
| Dh5α pBAD33::Nte5+Nti5 | Dh5α with pBAD33 encoding *Nte5+Nti5 under arabinose-inducible promoter control*; Cm<sup>R</sup> | This study |
| Dh5α pBAD33::Nte6<sup>R1300S</sup> | Dh5α with pBAD33 encoding *Nte6<sup>R1300S</sup> under arabinose-inducible promoter control*; Cm<sup>R</sup> | This study |
| Dh5α pBAD33::Nte6+Nti6 | Dh5α with pBAD33 encoding *Nte6+Nti6 under arabinose-inducible promoter control*; Cm<sup>R</sup> | This study |

**Table supplement 3.**
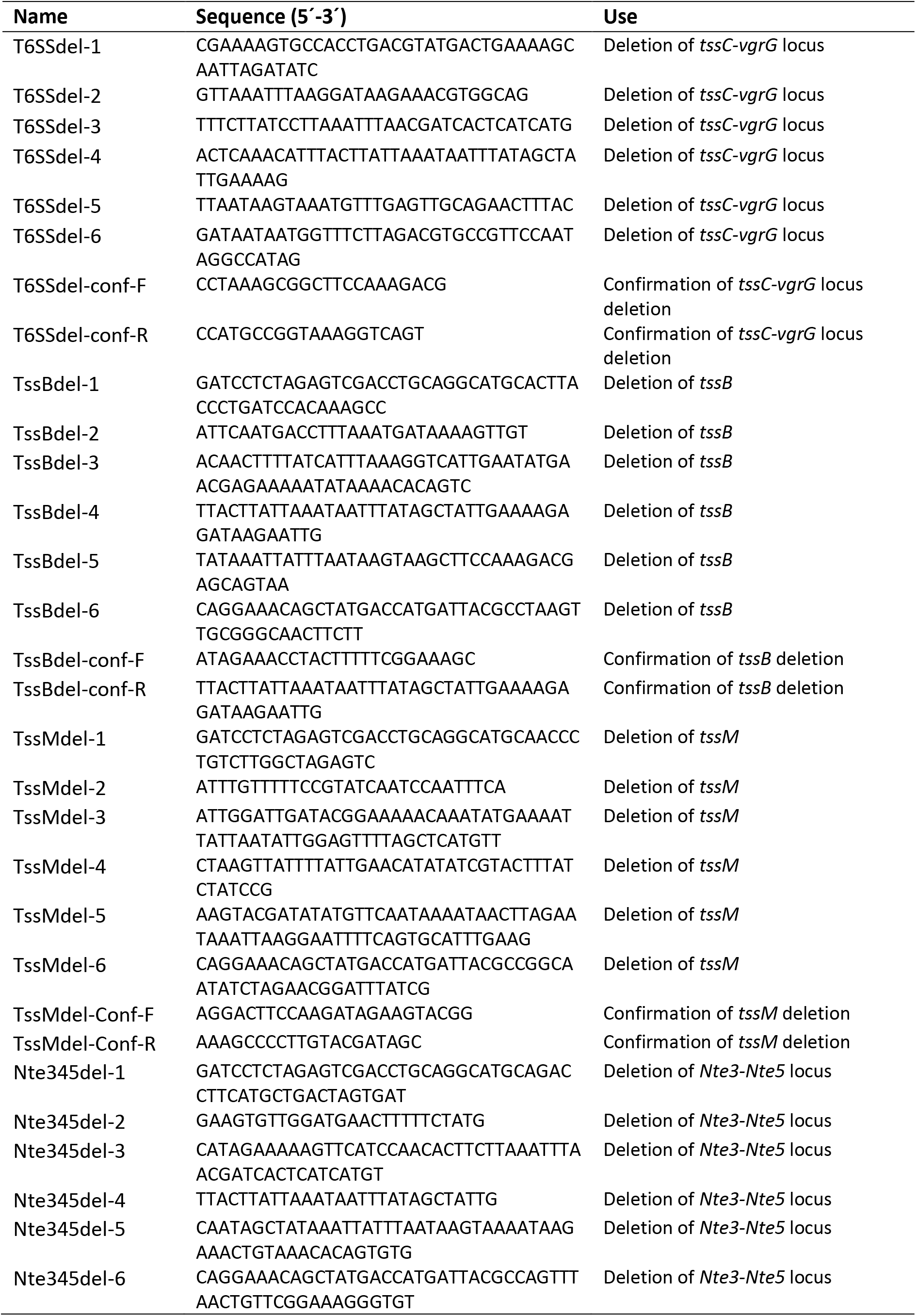
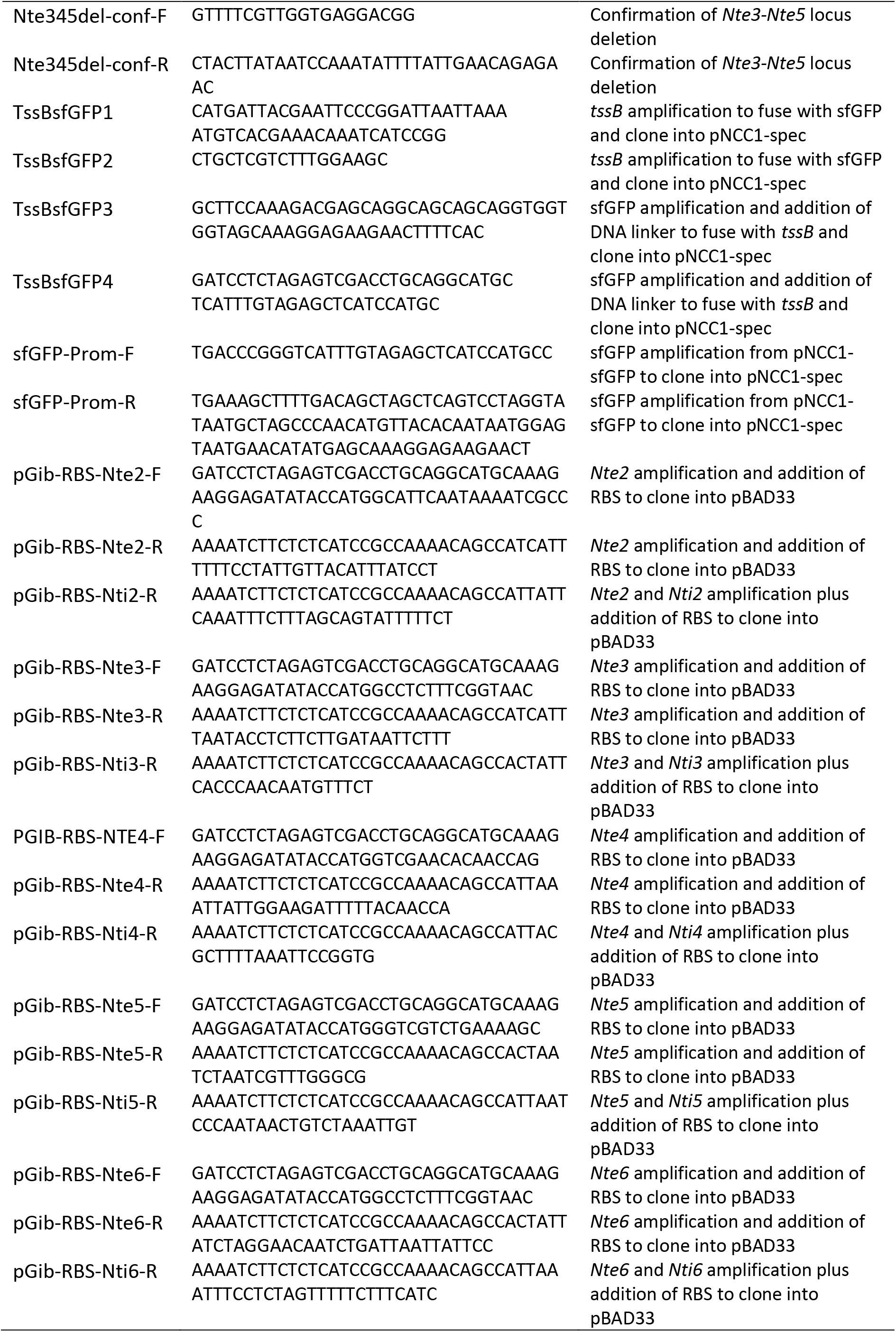
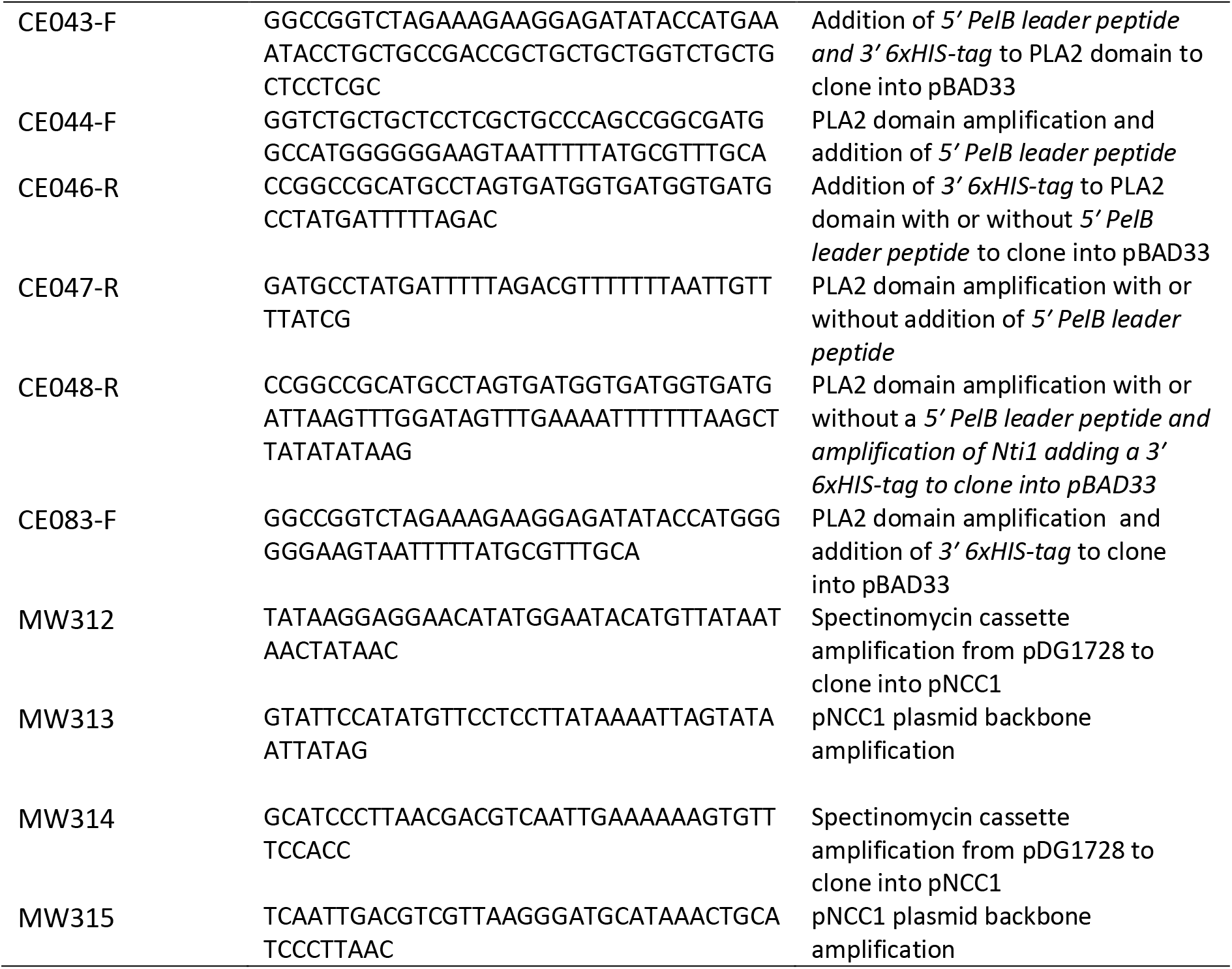
Primers used in this study.

## SUPPLEMENTARY MOVIES LEGENDS

**Movie supplement 1.** Visualisation of *N. cinerea* T6SS contraction.

*N. cinerea* 346TΔ*tssB::tssBsfGFP* (green) and prey cells *N. cinerea* 27178A_*sfCherry* (red) were mixed at a ratio of 1:1 and spotted on a 1% agarose PBS pad supplemented with 0.1 mM IPTG. The cells were imaged for 12 seconds with a rate of 1 image per second.

**Movie supplement 2.** Visualisation of *N. cinerea* T6SS foci.

*N. cinerea* 346TΔ*tssB*::*tssB*sfGFP (green) or *N. cinerea* 346T*ΔtssBΔtssM::tssBsfGFP* (green) and prey cells *N. cinerea* 27178A_*sfCherry* (red) were mixed at a ratio of 1:1 and spotted on a PBS 1% agarose pad supplemented with 0.1 mM IPTG. The cells were imaged for 200 seconds with a rate of one image per 10 seconds.

**Movie supplement 3.** *N. cinerea* T6SS elicits prey lysis.

*N. cinerea* 346TΔ*tssB*::*tssB*sfGFP (green) and *N. cinerea* 27178A_*sfCherry* (red) were mixed at a ratio of 1:1 and spotted onto agarose padS supplemented with 0.1 mM IPTG and the cell-impermeable DNA stain SYTOX Blue (0.5 μM) as an indicator for loss of membrane integrity. The cells were imaged for 10 min with an image taken every 10 seconds.

**Movie supplement 4.** Growing edge of colonies with a piliated attacker *N. cinerea* 346T*_gfp*, (green) and non-piliated prey 346TΔ*nte*/*i3*-*5*Δ*pilE1*/*2_sfCherry* (red).

Strains were spotted at a ratio of 1:1. Colonies were imaged every 2 h between 4 and 16 h post inoculation. Over time a population of non-piliated prey segregates to the edge, escape T6SS assault and dominates the growing colony.

**Movie supplement 5.** Growing edge of colonies with a piliated attacker *N. cinerea* 346T*_gfp*, (green) and piliated prey 346TΔ*nte*/*i3*-*5_sfCherry* (red).

Strains were spotted at a ratio of 1:1. Colonies were imaged every 2 h between 4 and 16 h post inoculation. Over time the T6SS expressing strain dominates the colony.

**Movie supplement 6.** Growing edge of colonies with two wild-type strains. *N. cinerea* 346T_*gfp* (green) and *N. cinerea* 346T_*sfCherry* (red).

Strains were spotted at a ratio of 1:1. Colonies were imaged between 4 and 16 h post inoculation. Images obtained every 2 h. No dominance of either strain is observed.

